# The conserved and divergent roles of Prdm3 and Prdm16 in zebrafish and mouse craniofacial development

**DOI:** 10.1101/844936

**Authors:** Lomeli Carpio Shull, Rwik Sen, Johannes Menzel, Kristin Bruk Artinger

**Affiliations:** Department of Craniofacial Biology, School of Dental Medicine, University of Colorado Anschutz Medical Campus, Aurora, CO 80045, USA; Cell Biology, Stem Cells, and Development Graduate Program, University of Colorado Anschutz Medical Campus, Aurora, CO 80045, USA; Molecular Biology Graduate Program, University of Colorado Anschutz Medical Campus, Aurora, CO 80045, USA

**Keywords:** Prdm3/Evi1/Mecom, Prdm16, neural crest, craniofacial, H3K9me3, H3K4me3

## Abstract

The formation of the craniofacial skeleton is a highly dynamic process that requires proper orchestration of various cellular processes in cranial neural crest cell (cNCC) development, including cell migration, proliferation, differentiation, polarity and cell death. Alterations that occur during cNCC development result in congenital birth defects and craniofacial abnormalities such as cleft lip with or without cleft palate. While the gene regulatory networks facilitating neural crest development have been extensively studied, the epigenetic mechanisms by which these pathways are activated or repressed in a temporal and spatially regulated manner remain largely unknown. Chromatin modifers can precisely modify gene expression through a variety of mechanisms including histone modifications such as methylation. Here, we investigated the role of two members of the PRDM (Positive regulatory domain) histone methyltransferase family, *Prdm3* and *Prdm16* in craniofacial development using genetic models in zebrafish and mice. Loss of *prdm3* or *prdm16* in zebrafish causes craniofacial defects including hypoplasia of the craniofacial cartilage elements, undefined posterior ceratobranchials, and decreased mineralization of the parasphenoid. In mice, while conditional loss of *Prdm3* in the early embryo proper causes mid-gestation lethality, loss of *Prdm16* caused craniofacial defects including anterior mandibular hypoplasia, clefting in the secondary palate and severe middle ear defects. In zebrafish, *prdm3* and *prdm16* compensate for each other as well as a third Prdm family member, *prdm1a.* Combinatorial loss of *prdm1a*, *prdm3*, and *prdm16* alleles results in severe hypoplasia of the anterior cartilage elements, abnormal formation of the jaw joint, complete loss of the posterior ceratobranchials, and clefting of the ethmoid plate. We further determined that loss of *prdm3* and *prdm16* reduces methylation of histone 3 lysine 9 (repression) and histone 3 lysine 4 (activation) in zebrafish. In mice, loss of *Prdm16* significantly decreased histone 3 lysine 9 methylation in the palatal shelves but surprisingly did not change histone 3 lysine 4 methylation. Taken together, *Prdm3* and *Prdm16* play an important role in craniofacial development by maintaining temporal and spatial regulation of gene regulatory networks necessary for proper cNCC development and these functions are both conserved and divergent across vertebrates.

## Introduction

The craniofacial skeleton forms through tightly regulated cellular processes involving extensive cross-talk between multiple cell types and tissues [1]. During development, cranial neural crest cells (cNCCs), a multipotent stem-cell like population of cells, arise at the neural plate border and undergo an epithelial-to-mesenchymal transition, which activates their migration toward the pharyngeal arches [2, 3]. Once cNCCs have populated the pharyngeal arches, they initiate differentiation programs toward chondrocyte and osteoblast lineages to form the cartilage and bones of the face, among other cranial tissue derivatives such as the cranial ganglia of the peripheral nervous system (PNS) [2, 3]. NCCs that populate the pharyngeal arches retain a precise medial-lateral and anterior-posterior segmental identity from their position of origin. Cells from the midbrain and hindbrain rhombomere (r) 1 and 2 populate pharyngeal arch (PA)1, cells from r4 migrate to PA2 and cells from r6-8 migrate into posterior PA3-7 [4–9]. Once in the arches, cNCCs engage in reciprocal interactions with both surrounding cells and the environment. The developmental pathways crucial for the proper formation of the craniofacial skeleton need to by tightly regulated. Regulatory cascades such as Hox genes, including the Hoxa paralogs, are key for patterning within the posterior pharyngeal arches which is key for positioning the posterior cartilage elements including thyroid cartilage, cricoid cartilages and tracheal rings [6, 10, 11]. Perturbations to these developmental processes leads to birth defects, including craniofacial abnormalities such as cleft lip with or without cleft palate [12]. While the gene regulatory networks (GRNs) and pathways driving cNCC development have been identified and extensively studied, the mechanisms by which these networks are orchestrated spatially and temporally remain largely unknown.

Chromatin modifiers control gene expression epigenetically by regulating chromatin assembly and compaction through a variety of different mechanisms including DNA methylation, histone modification, nucleosome positioning, chromatin dynamics and chromatin interactions [13, 14]. Mutations in chromatin modifiers or disruption of their enzymatic activity can influence the expression of downstream target genes and alter associated GRNs that control important biological processes. The role of epigenetic regulators during cNCC development has recently become more appreciated, due to the identification of histone modifying enzymes associated with craniofacial phenotypes. Among these are MLL2 and KDM6A in Kabuki syndrome and CHD7 in CHARGE syndrome [15]. We have added the PRDM (Positive Regulatory Domain) family of lysine methyltransferases to the list, which control gene expression by modifying chromatin at specific target gene promoters [16–18]. PRDMs are characterized by the presence of a PR domain which shares homology to the SET domain found in many histone lysine methyltransferases, as well as having a variable number of zinc finger domains [17]. Many PRDMs are involved in specific cellular processes including differentiation and cell fate determination. While all PRDMs carry the SET domain, few have intrinsic methyltransferase activity. Instead, many PRDMs govern gene expression by interacting in complexes with other histone modifying proteins to elicit their role in epigenetic gene regulation [18]. Human genome-wide association studies have linked the genes encoding two PRDMs, PRDM3 (EVI1/MECOM) and PRDM16, to cleft palate [19, 20]. Human mutations in PRDM1 (Blimp1) have been linked to Split/Hand Foot malformation, a disorder that causes severe limb deformities and varying degrees of craniofacial abnormalities (Artinger lab, unpublished) [17]. The specific role of PRDM3 and PRDM16, and their genetic interaction with PRDM1 during cNCC development remains largely unknown.

Unlike most PRDM family members, PRDM3 and PRDM16 exhibit intrinsic methyltransferase activity and both modify histone H3 lysine 9 (H3K9), a mark associated with transcriptional repression [21]. Specifically, PRDM3 and PRDM16, *in vitro*, mediate a stepwise process of H3K9me monomethylation to H3K9 trimethylation to establish heterochromatin stability and transcriptional repression [21]. Other studies have implicated PRDM3 and PRDM16 as activators of gene expression *in vitro* through methylation of histone 3 lysine 4 (H3K4) [21–24]. PRDM1 has no known intrinsic methyltransferase activity *in vivo.* Instead, PRDM1 forms complexes with other co-factors and histone-modifying enzymes including the methyltransferase, G9a [25]. PRDM3 and PRDM16 can also indirectly regulate gene expression by forming complexes with other co-factors and enzymes like histone acetyltransferases and histone deacetylases [17, 26–30]. Moreover, these three PRDMs contain putative DNA binding sites, and they also possess zinc finger domains, implicating their potential to directly bind DNA to also mediate gene expression [16–18, 31–34].

PRDM1, PRDM3 and PRDM16 are important in a variety of developmental processes including early embryogenesis, limb, hematopoiesis, plasma cell differentiation, brown adipocyte differentiation, and formation of the cerebral cortex [23, 34–50]. Loss of *Prdm1* in mice leads to defects in posterior pharyngeal arch development and analysis of a null allele in zebrafish results in loss of posterior ceratobranchial cartilages, and hypoplasia of the anterior cartilage elements [51, 52]. Individual knockdown or combinatorial knockdown of *prdm1a*, *prdm3* and *prdm16* in zebrafish causes craniofacial defects including hypoplasia of cartilage elements [36]. In mice, while *Prdm3* null mice die early during embryogenesis, they present with hypoplastic pharyngeal arches [53]. Loss of *Prdm16* in a recessive ENU-derived mutant causes cleft palate, mandibular hypoplasia, and defects in middle ear formation [35, 54]. While this evidence suggests the importance of *prdm3* and *prdm16* in craniofacial development, the exact mechanism of their action, and interaction together with *prdm1a*, in mediating proper cNCC development and the proper formation of the craniofacial skeleton has not been studied in detail, especially in mammals.

Here, we examined the role of *Prdm3* and *Prdm16* during craniofacial development using genetic models in zebrafish and mice. Loss of *prdm3* or *prdm16* in zebrafish causes craniofacial defects including hypoplasia of the craniofacial cartilage elements, undefined posterior ceratobranchials, and decreased mineralization of the parasphenoid. In mice, loss of *Prdm3* in the early embryo proper with the *Sox2-Cre* driver causes mid-gestation lethality. Conversely, conditional loss of *Prdm16* in the embryo epiblast was not lethal but instead resulted in craniofacial defects including anterior mandibular hypoplasia, and severe hypoplastic tympanic rings and middle ear defects. A subset of these mutants develop a secondary cleft palate. In zebrafish, we show genetic compensation between *prdm3, prdm16* as well as a third Prdm family member, *prdm1a.* Combinatorial loss of different *prdm1a*, *prdm3*, *prdm16* alleles causes more severe craniofacial phenotypes in zebrafish, including severe hypoplasia of the anterior cartilage elements, partial loss of the jaw joint with fusion of the Meckel’s cartilage to the palatoquadrate, complete loss of the posterior ceratobranchials, and clefting of the ethmoid plate. Finally, loss of *prdm3* and *prdm16* reduces methylation of their target substrates, histone 3 lysine 9 (repression) and histone 3 lysine 4 (activation) H3K4 in zebrafish. In mice, loss of *Prdm16* significantly decreased histone 3 lysine 9 methylation in the palatal shelves but surprisingly did not change histone 3 lysine 4 methylation. These results suggest loss of Prdm3 and Prdm16 in zebrafish and mice lead to altered chromatin structure which disrupts downstream gene expression profiles. All together, these results demonstrate that *Prdm3* and *Prdm16* play an important role in craniofacial skeletal formation by maintaining temporal and spatial regulation of gene expression and downstream signaling factors that are necessary for proper cNCC development and craniofacial formation and these functions are conserved across vertebrates.

## Materials and Methods

### Animals

Zebrafish were maintained as previously described [55]. Embryos were raised in defined Embryo Medium at 28.5°C and staged developmentally following published standards as described [56]. The wildtype (WT) strain used include the AB line (ZIRC). The Institutional Animal Care and Use Committee of the University of Colorado Anschutz Medical Campus approved all animal experiments performed in this study and conform to NIH regulatory standards of care.

### Mice

*Prdm3^fl/fl^* [44]*, Prdm16^fl/fl^* (Jackson Laboratories), and *Tg(Sox2-Cre)* [57] mice were maintained on the C57/Bl6 background. For timed matings, *Prdm3^fl/fl^*or *Prdm16^fl/fl^* females were bred to *Prdm3^fl/+^;Sox2-Cre^+/Tg^*(also denoted as *Prdm3^Δ/+^;Sox2-Cre^+/Tg^*) or *Prdm16^fl/+^;Sox2-Cre^+/Tg^*(also *Prdm16^Δ/+^;Sox2-Cre^+/Tg^*) males, respectively. The morning a vaginal plug was detected was signified as embryonic day 0.5. Embryos of matching somite numbers were used for experiments. Mice were bred and maintained in accordance with the recommendations in the Guide for the Care and Use of Laboratory Animals of the National Institutes of Health. The protocol was approved by the Institutional Animal Care and Use Committee of the University of Colorado Anschutz Medical Campus.

### Generation of Zebrafish Mutant Lines

Zebrafish mutant lines for *prdm3* and *prdm16* were generated via CRISPR-based mutagenesis. CRISPR target sites were identified using the ZiFiT Targeter (http://zifit.partners.org/ZiFiT/). Selected targets were used to generate guide RNAs (gRNAs) specific for *prdm3* or *prdm16*. gRNAs were prepared as previously described [58]. Prepared gRNAs were then co-injected with 600 ng/ul of Cas9 Protein (PNA bio) and 200 mM KCl. Cas9/sgRNA complexes were formed by incubating Cas9 protein with sgRNA at room temperature for 5 minutes prior to injecting into the cytoplasm of WT AB zebrafish embryos at the 1-cell stage. Mosaic germline founders were identified by screening the progeny of CRISPR injected fish grown to adulthood. Two separate alleles were identified for each single mutant gene, *prdm3^CO1005^*, *prdm16^CO1006^*, *prdm3^CO1013^*, and *prdm16^CO1014^*. For this study, we focused on *prdm3^CO1005^* and *prdm16^CO1006^*alleles schematized in Supplemental Figure 1. From here on, these alleles will be referred to as *prdm3-/-* for *prdm3^CO1005^* and *prdm16-/-* for *prdm16^CO1006^*. All *prdm3* and *prdm16* mutant alleles are predicted frameshift mutations, leading to reduced open reading frames that interrupt the coding sequence upstream of the PR/SET domain responsible for functional histone methyltransferase activity.

### Genotyping

For fish, fin clips, single whole embryos or single embryo tails were lysed in Lysis Buffer (10 mM Tris-HCl (pH 8.0), 50 mM KCl, 0.3% Tween-20, 0.3% NP-40, 1 mM EDTA) for 10 minutes at 95° C, incubated with 50 ug of Proteinase K at 55°C for 2 hours, followed by another incubation at 95°C for 10 minutes. For genotyping *prdm3* mutants, the following primers were used (F) 5’-CTGTGGGCAGATGTTTAGCA-3’ and (R) 5’-ACTATGATGCCGGTTTGTCC-3’. The 200 bp PCR product was run out on a high percentage (4%) agarose gel to distinguish homozygous mutant embryos from heterozygous and wildtype embryos. *prdm16* CRISPR mutants were genotyped using the following primer sets (F) 5’-CAGATGACAGCGAGGCAGTA-3’ and (R) 5’-CGTCGCACTCTGCAGTCCGTTGTATGG-3’. This 300 bp PCR product was diluted 1:100 in nuclease-free water. A second set of primers was designed using dCAPS Finder 2.0 (http://helix.wustl.edu/dcaps/) to generate a primer containing a mismatch that creates a restriction endonuclease site based on the identified SNP mutation in the *prdm16* mutant allele. This primer set, (F) 5’-GGGTCCCGCATCCCCTCAGCCCTGCCT-3’ and (R) 5’-CGTCGCACTCTGCAGTCCGTTGTATGG-3’, generates a 200 bp PCR product that was then digested using the restriction enzyme, XhoI, which digests the wildtype sequence but is unable to cut the mutant sequence, allowing for homozygous mutant embryos to be distinguished from heterozygous and wildtype embryos.

For mice, tail clips from weanlings and tail clips or yolk sacs from embryos were lysed in DNA Lysis Buffer (10 mM Tris-HCl (pH 8.0), 100 mM NaCl, 10 mM EDTA (pH 8.0), 0.5% SDS) and 100 ug of Proteinase K overnight at 55°C. Genomic DNA was isolated following phenol/chloroform extraction. DNA pellets were air dried and resuspended in nuclease-free water. For *Prdm3* genotyping, the following primer sets were used to detect the floxed allele, (F) 5’-GCATTGCTTCATTTAGATCCC-3’ and (R) 5’-CAGGGAATTATGACTTTGAGGAAA-3’ and a second reverse primer (R) 5’-TGGTCAAAGCTTTCAGGGTT-3’ was used in combination with the same forward primer to detect the deleted allele. For *Prdm16* genotyping, the following primer sets were used: (G1) 5’-TGCTAAGCCTTCACCGTTCT-3’, (G2) 5’-TGCAGGGAGATTGACAAGTG-3’ and (G3) 5’-CCATGGTTCACATGGTCAAG-3’. This primer set detected the floxed allele at 490 bp, the wildtype allele at 408 bp and the deleted allele at 244 bp. For *Sox2-Cre* PCR genotyping, the following primer set was used: (Transgene F) 5’-CTCTAGAGCCTCTGCTAACC-3’, (Transgene R) 5’-ACATGTCCATCAGGTTCTTGC-3’, (Internal Positive Control F) 5’-CTAGGCCACAGAATTGAAAGATCT-3’, (Internal Positive Control R) 5’-GTAGGTGGAAATTCTAGCATCATCC-3’, where the transgene was detected at 250 bp and the internal positive control was detected at 324 bp.

### Skeletal Staining

For zebrafish, Alcian Blue (cartilage) and Alizarin Red (bone) staining was performed at room temperature as previously described [59]. Briefly, larvae were anesthetized in Tricaine (Sigma A5040) and fixed for one hour in 2% PFA. Following a 10-minute rinse in 100 mM Tris (pH 7.5) and 10 mM MgCl_2_, larvae were incubated in Alcian Blue solution (0.04% Alcian Blue, 80% ethanol, 100 mM Tris (pH 7.5), 10 mM MgCl_2_) overnight at room temperature. Larvae were then rehydrated and destained through a gradient of ethanol solutions 80%, 50%, 25% ethanol containing 100 mM Tris (pH 7.5) and 10 mM MgCl_2_, then bleached for 10 minutes in 3% H_2_O_2_ with 0.55% KOH at room temperature, washed twice in 25% glycerol with 0.1% KOH, then stained in Alizarin Red (0.01% Alizarin Red dissolved in 25% glycerol and 100 mM Tris (pH 7.5)) for 30 to 45 minutes at room temperature. Samples were destained in 50% glycerol with 0.1% KOH. Whole-mount and dissected and flat mounted specimens were mounted in 50% glycerol and imaged with LAS v4.4 software on Leica.

For mice, Alcian blue and Alizarin red staining was performed as previously described [60] for E18.5 embryos. Briefly, mouse embryos were harvested at E18.5 in 1xPBS. Skin and internal organs were removed. Samples were fixed overnight in 95% ethanol at room temperature followed by an incubation in 100% acetone for 2 days at room temperature. Embryos were stained in Alcian Blue/Alizarin Red staining solution (0.015% Alcian Blue, 0.05% Alizarin Red, 5% Glacial acetic acid in 70% ethanol) for 3 days at 37°C then rinsed one time in water before being cleared in 1% KOH overnight at room temperature. Embryos were then transferred through a series of solutions with decreasing KOH concentrations and increasing glycerol concentrations. Stained skeletons were stored and imaged in 80% glycerol on Leica stereomicroscope with the LAS v4.4 software.

For mouse embryos staged to E13.5, embryos were harvested in 1x PBS and fixed overnight in 95% ethanol. Embryos were incubated in 70% ethanol, then washed in 5% acetic acid for two hours at room temperature. Embryos were then incubated in Alcian Blue staining solution (1% acetic acid, 0.015% Alcian blue) for 3 days at room temperature followed by incubation for 2 hours in 5% acetic acid, then 2 hours in 100% methanol at room temperature. Embryos were then destained and stored in benzyl benzoate:benzyl alcohol solution before being imaged on Leica steromicroscope with the lAS v4.4 software.

### Histology

Embryos were collected at E15.5 in PBS. The heads of the animals were removed and fixed in 4% paraformaldehyde overnight then dehydrated through a graded ethanol series, embedded in paraffin and sectioned to a thickness of 8 µm onto glass slides. After deparaffinization and rehydration, sections were stained with Weigert’s iron hematoxylin, 0.05% Fast Green, and 0.1% Safranin O and mounted with Permount (Electron Microscopy Sciences).

### In situ hybridization

Zebrafish embryos were collected at 48 hours post fertilization and fixed in 4% paraformaldehyde overnight and dehydrated in methanol. Following rehydration in methanol gradients, embryos were bleached (3% hydrogen peroxide and 0.5% potassium hydroxide) until body pigment cells and the eyes were visibly cleared (10 minutes), digested with proteinase K for 8 minutes and fixed in 4% paraformaldehyde for 25 mins at 4°C. Embryos and DIG-labeled probes (*col2a1, sox9a, dlx2a, crestin*) were prehybridized for 2-4 hours at 65°C before incubating embryos with the indicated probes overnight at 65°C. Embryos were subjected to a series of stringency washes in a wash buffer containing 50% formamide, 2X SSC pH 4.5, and 1% SDS before blocking in a buffer containing 2% Blocking Reagent (Roche) and 20% Heat-Inactivated sheep serum diluted in 1X MABT (100 mM maleic acid, 150 mM NaCl, 0.1% Tween-20) and anti-DIG antibody (Roche) incubation overnight at 4°C. After another set of stringency washes in NTMT (100 mM NaCl, 100 mM Tris pH 9.5, 50 mM MgCl_2_, 1% Tween-20), embryos were developed with BM purple reagent (Roche) and stored in PBS.

### RNA isolation and qPCR

Total RNA was isolated from pooled heads of *prdm3-/-*, *prdm16-/-* and wildtype 48 hpf embryos with TRIzol reagent (Invitrogen) and phenol/chloroform. RNA (1-1.5 μg) was reverse transcribed to complementary DNA (cDNA) with SuperScript III First-Strand Synthesis cDNA kit (Invitrogen) for real-time semiquantitative PCR (qPCR) with primers for *prdm1a* (TaqMan Probe Assay), *prdm3* ((F) 5’-CACCCAAGACTGACCCATCC-3’, (R) 5’-GAAGTGTAGTGGGTGGAGCC-3’), *prdm16* ((F) 5’-GCCCCAAAGCTTTCAACTGG-3’, (R) 5’-CTGGGGTCTGTGAACACACTA-3’) and SYBR Green master mix (Biorad) or TaqMan master mix for *prdm1a.* Transcript levels were normalized to reference gene, *rpl13a*. Transcript abundance and relative fold changes in gene expression were quantified using the 2^-ΔΔCt^ method relative to control.

### Western Blot

For western blotting, 30 to 40 zebrafish embryos of each genotype were collected and pooled at 48 hpf. Embryos were incubated on ice for 5 minutes. Calcium-free Ginzberg Fish Ringer’s solution (NaCl, KCl, NaHCO3) was added to the embryos for deyolking. Samples were centrifuged for 1.5 minutes at 5000 rpm. Pellets were washed in deyolking wash buffer (NaCl, Tris (pH 8.5), KCL and CaCl_2_). Samples were again centrifuged for 1.5 minutes at 5000 rpm. All liquid was removed, and pellet was resuspended and lysed in SDS sample buffer (0.1% glycerol, 0.01% SDS, 0.1 M Tris (pH 6.8)) for at least 10 minutes on ice. Embryos were homogenized in lysis buffer and briefly centrifuged. Total protein concentrations were determined with the Bio-Rad Dc Protein assay. Proteins (20 ug) were separated by SDS-PAGE (12%) and transferred to polyvinylidene difluoride membranes. Membranes were blotted using antibodies for H3K9me3 (Abcam, ab8898), H3K4me3 (Cell Signaling Technology, 9751) and H3 (Cell Signaling Technology, 9715S) and corresponding secondary antibodies. Chemiluminescent detection was performed with Luminata Classico Western HRP Substrate (Millipore) and imaged on a BioRad Chemidoc multiplex imager. Band intensity was quantified in ImageJ. For each replicate experiment, the band intensity for each sample for each histone mark (H3K9me3 or H3K4me3) was normalized to that sample’s respective Total H3 loading control. The normalized band intensities for all wt/het controls across experimental replicates was then averaged. Normalized (by Total H3) band intensities for each histone mark for the the mutant lysate samples were then normalized to the averaged wt/het controls and then averaged across experimental replicates.

### Immunofluorescence

For immunofluorescence on coronal mouse sections, sections were deparaffinized and rehydrated then subjected to antigen retrieval using a ready-to-use Citrate target retrieval solution, pH 9.5 (Dako, S1700) for 20 minutes at 90°C. Sections were cooled before washing 2 times briefly in Tris-Buffered Saline (TBS) with 0.025% Triton X-100. Sections were then blocked in 10% normal goat serum with 1% Bovine Serum Albumin in TBS for 2 hours at room temperature before the primary antibody was added. H3K9me3 (Abcam, ab8898) or H3K4me3 (Cell Signaling Technology, 9751) primary antibodies were diluted in blocking solution and incubated on sections overnight at 4°C. Slides were rinsed 2 times briefly in TBS with 0.025% Triton X-100 then incubated in appropriate goat anti-rabbit secondary antibodies diluted in blocking solution for 1 hour at room temperature. Sections were then washed in TBS with 0.025% Triton X-100 briefly followed by a quick 5 minute incubation in TBS with DAPI and another quick rinse in TBS with 0.025% Triton X-100 before mounting with Vectashield. Sections were imaged on a Leica TCS SP8 confocal microscope and images were processed with the LASX software. Image quantification was carried out using ImageJ.

### Whole Mount Immunofluorescence

Zebrafish embryos were collected at the indicated timepoint and fixed in 4% paraformaldehyde overnight at 4°C, embryos were washed and dehydrated through a graded methanol series. Samples in 100% methanol were incubated at least overnight in methanol prior to the next steps. Following rehydration through the reverse methanol gradient, embryos were washed in 1x Phosphate Buffered Saline (PBS, pH 7.3) with 0.1% Tween 3 times for 10 minutes at room temperature, then equilibrated in 150 mM Tris pH 9.5 for 5 minutes at room temperature before a 20 minute incubation in the Tris solution at 70°C. Following this retrieval step, embryos were washed 3 times for 10 minutes each in PBS with 0.1% Tween at room temperature then 2 times for 5 minutes in distilled water before blocking at room temperature in blocking solution containing 10% normal goat serum and 1% bovine serum albumin in PBS. Embryos were then incubated in primary antibodies for 3 days at 4°C. Primary antibodies, phosphoH3 (Sigma, H0412) and cleaved caspase 3 (Cell Signaling, 9661) were diluted in blocking solution. Following primary antibody incubation, samples were washed thoroughly in PBS with 0.1% Triton X-100 then incubated in fluorescently tagged goat anti-rabbit 546 secondary antibodies (Invitrogen) for 2 days at 4°C. After secondary antibody incubation, embryos were washed thoroughly in PBS with 0.1% Triton X-100 before incubating in DAPI diluted in PBS for 1 hour at room temperature. Samples were quickly washed in PBS and passed through a glycerol/PBS gradient before mounting on glass slides. Embryos were imaged on a Leica TCS SP8 confocal microscope. Images were processed in LASX software and quantification of cell numbers were completed on max projections of z-stack images using Image J.

### Statistical Analysis

Data shown are the means ± SEM from the number of samples or experiments indicated in the figure legends. All assays were repeated at least three times with independent samples. *P* values were determined with Student’s *t* tests.

## Results

### Deletion of *prdm3* causes craniofacial defects in zebrafish and mid-gestation lethality in mice

Previous studies have shown *prdm3* and *prdm16* are expressed in the pharyngeal arches and the head folds during zebrafish and mouse development, respectively [36, 41, 61]. Knockdown of *prdm3* and *prdm16* by morpholino disrupts formation of the craniofacial skeleton [36]. To better understand the mechanisms of *prdm3* and *prdm16* in cranial neural crest cells and the development of the craniofacial skeleton, we utilized CRISPR/Cas9 to generate loss-of-function alleles. Guide RNAs were designed to target the PR/SET domain, induce early frameshift mutations, and abrogate protein function (Supplemental Fig. 1A-1B).

To assess the consequences on the development of the craniofacial skeleton with loss of *prdm3*, mutant zebrafish embryos were collected at 6 days post fertilization (dpf) and stained with Alcian blue and Alizarin red to evaluate cartilage and bone development. *prdm3-/-* mutants were present in Mendelian ratios and showed mild craniofacial defects (Fig. 1A). The defects present in dissected and mounted viscerocraium and neurocranium tissues included mild hypoplasia of Meckel’s cartilage and palatoquadrate, and undefined posterior ceratobranchial cartilages (Fig. 1A-1F). Specifically, there was a 20% reduction in the distance measured from the Meckel’s cartilage to the beginning of the palatoquadrate (Fig. 1A). In the neurocranium, *prdm3* mutants have a 13% reduction in the distance between trabeculae (Fig. 1A, 1D) and a 53% decrease in length of the parasphenoid (Fig. 1E), a neural crest derived tissue, that along with the ethmoid plate is considered equivalent to mammalian palatal tissue [62]. In addition, the area of the mineralized tissue of the parasphenoid is reduced by 31%. (Fig. 1A, 1F).

**Figure 1.**
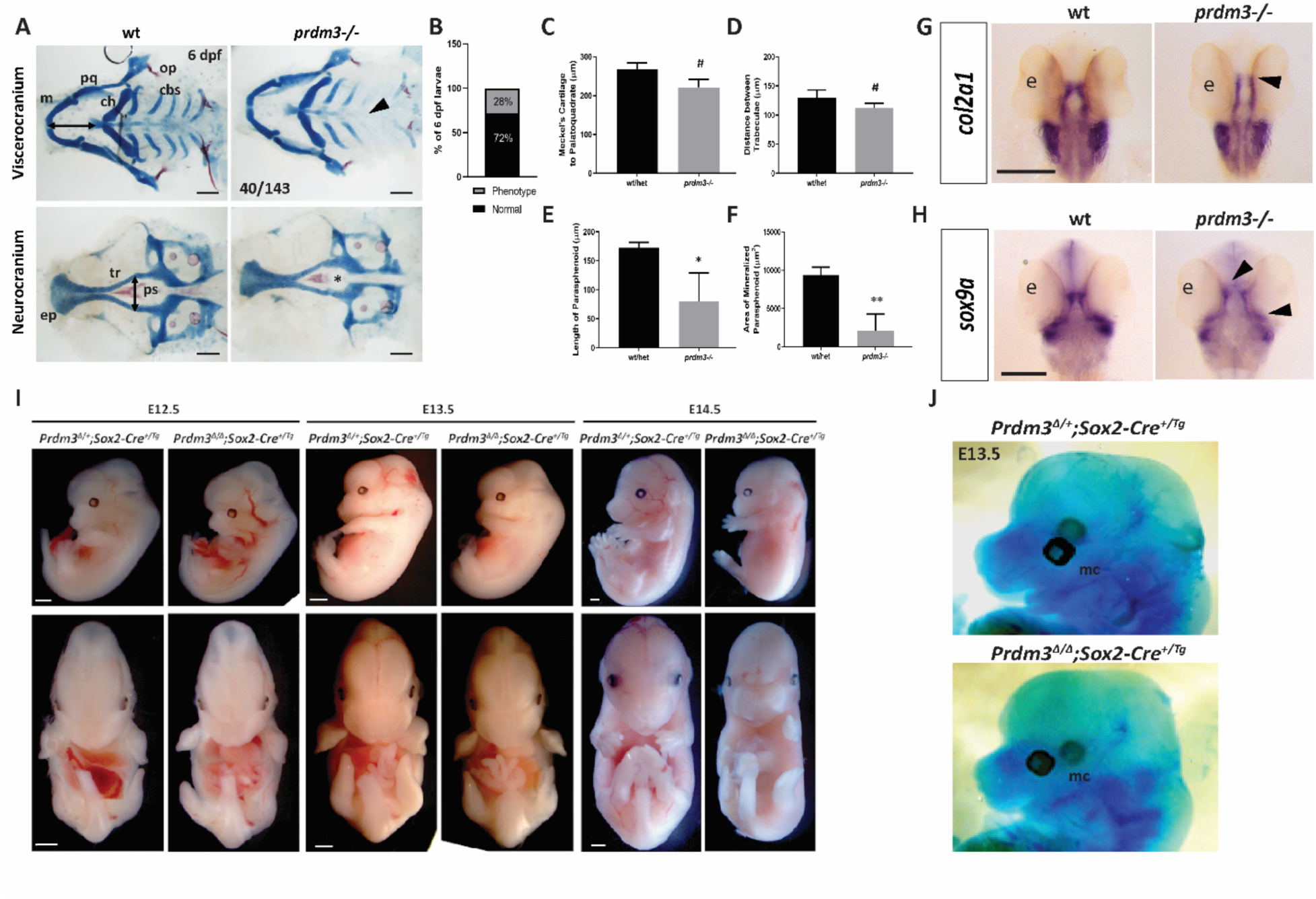
Genetic ablation of PRDM3 causes subtle craniofacial defects in zebrafish and mid-gestation lethality in mice. (A-E) Wildtype and *prdm3-/-* zebrafish embryos were collected at 6 dpf and stained for Alcian blue and Alizarin red. (A) Images of dissected viscerocranium and neurocranium of wildtype (wt) and *prdm3-/-* embryos. *prdm3-/-* embryos and their cartilage phenotypes were present in Mendelian ratios (B). (C-F) Quantification of cartilage elements, Meckel’s cartilage to palatoquadrate (horizontal black double arrow in wt viscerocranium of A) (C), distance between trabeculae (vertical black double arrow in wt neurocraium of A) (D), length of parashenoid tissue (E), and area of mineralized parasphenoid (F). ceratobranchial arches (cbs), ceratohyal (ch), Meckel’s cartilage (m), palatoquadrate (pq), parasphenoid (ps), trabeculae (tr), (n = 6 for each group measured). Scale bar, 100 µm. (G-H) Ventral views of *in situ hybridization* for *col2a1* (G) and *sox9a* (H) in *prdm3* mutants or wildtype controls at 48 hpf. Black arrow heads indicate decreased expression in the cartilage elements of the neurocranium and viscerocranium. Scale bar, 250 µm. (I-J) *Prdm3^fl/fl^*females were bred for timed matings to *Prdm3^Δ/+^;Sox2^+/Tg^*males and embryos were collected at the indicated timepoints. Shown are lateral (top) and ventral (bottom) gross phenotypes of mutant animals (I). Scale bar 750 µm. (J) *Prdm3^Δ/Δ^;Sox2^+/Tg^* and control embryos were collected at E13.5 and stained with Alcian blue. Shown are lateral views of the head. Meckel’s cartilage (mc). * p ≤ 0.05, ** p ≤ 0.005, # p ≤ 0.1, Student’s *t* test.

To further understand the mild craniofacial phenotypes of *prdm3*-deficient embryos, *in situ hybridization* for cartilage markers, type 2 collagen (*col2a1*) and SRY-box transcription factor (*sox9a*), were performed at 48 hpf (Fig. 1G-1H). Loss of *prdm3* lowered the expression of both *col2a1* and *sox9a* in the pharyngeal skeleton, suggesting changes to chondrocyte matrix proteins during craniofacial development and overall loss of chondrocyte functionality. There were no changes in levels of expression or localization of neural crest cell makers, *dlx2a* or *crestin* at 24 hpf (Supplemental Fig. 2A, 2B). Additionally, immunostaining for phosphorylated histone H3 or cleaved caspase 3 revealed no change in proliferation or cell death, respectively, in pre-chondrocytes of the pharyngeal arches at 24 hpf (Supplemental Fig. 3A-3D). These results suggest that loss of *prdm3* does not impede neural crest specification or migration. Neural crest cells are able to reach the pharyngeal arches and there is no reduction in cell proliferation or increase in cell death within these tissue regions. These results instead support the role for *prdm3* in the differentiation of neural crest cells.

To determine if PRDM3 function is conserved in mammals, we conditionally deleted *Prdm3* (*Evi1/Mecom*) in the early mouse embryonic epiblast using the *Sox2-Cre* driver (Fig. 1I) to circumvent the placental defects in Prdm3 knock-out embryos. The *Prdm3^Δ/Δ^;Sox2-Cre^+/Tg^* mutant animals did not have any noticeable placental defects but the *Prdm3* mutants failed to develop normally after E13.5. While we did not thoroughly investigate the cause of death, early lethality likely resulted from vascular abnormalities including visible defects in vasculogenesis and hemorrhaging, leading to embryonic lethality detected around E13.5-E14.5 (Fig 1I) and subsequent embryonic tissue breakdown and resorption at E14.5. These results are consistent with previous studies in mice that show disruption of *Prdm3/Evi1* causes lethality at E10.5 from cardiovascular and/or placental defects [53].

The lethality with early embryonic deletion of *Prdm3* hinders the ability to assess if craniofacial skeleton formation is initiated. To determine if craniofacial cartilaginous elements, namely Meckel’s cartilage, are at least present in *Prdm3^Δ/Δ^;Sox2-Cre^+/Tg^*mutants, E13.5 embryos were collected and stained for Alcian blue. Whole mount skeletal preparations showed that the craniofacial skeletal elements, specifically the Meckel’s cartilage forms but appears thicker and does not extend out as far anteriorly when compared to control littermates (Fig 1J). Noticeably, the head size of *Prdm3^Δ/Δ^;Sox2-Cre^+/Tg^*mutants was reduced compared to control animals, but this is proportionate to an overall reduction in embryo body size.

### Loss of *prdm16* in zebrafish and mice causes craniofacial abnormalities

To understand the role of PRDM16 in cranial neural crest cell development and the formation of the craniofacial skeleton, *prdm16-/-* zebrafish embryos were collected at 6 dpf and cartilage and bone were stained with Alcian blue and Alizarin red, respectively. Dissected viscerocranium and neurocranium tissues showed *prdm16-/-* embryos had a moderately more dramatic craniofacial phenotype compared to *prdm3-/-* embryos (Fig. 2A-2F). *prdm16-/-* mutants and their craniofacial skeleton phenotypes were present at Mendelian ratios (Fig. 2B). In the viscerocranium, *prdm16* mutants developed hypoplasia of the anterior cartilage elements, i.e., Meckel’s cartilage and the palatoquadrate, including a 23% reduction in the distance of the Meckel’s cartilage to the palatoquadrate as well as a more dramatic loss of the posterior ceratobranchial cartilages (Fig. 2A, 2C), relative to *prdm3* mutants (Fig. 1A-F). In the neurocranium, *prdm16-/-* zebrafish had a 11% decrease in the distance between the trabeculae (Fig. 2D), as well as 51% reduction in length of the parasphenoid (Fig. 2E) and a 31% decrease in the area of parasphenoid mineralized tissue (Fig. 2F). *In situ hybridization* for *col2a1* and *sox9a* showed reduced expression in the developing pharyngeal skeleton at 48 hpf in *prdm16-/-* compared to wildtype controls (Fig. 2G-2H). We noted subtle reductions in expression of *dlx2a* or *crestin* at 24 hpf (Supplemental Fig. 4A, 4B), particularly with a decrease in *dlx2a* in posterior arch 4 at 24 hpf (Supplemental Fig. 4A) and a decrease in *crestin* expression in the more anterior aches (Supplemental Fig. 4B). While there are minor changes in the expression of these markers, the neural crest is still specified in *prdm16* mutants. Neural crest cells are also still able to migrate and populate the pharyngeal arches with loss of *prdm16*. Proliferation and cell death, as assessed by immunofluorescence staining for phosphorylated histone H3 and cleaved caspase 3, respectively, were unchanged in the pharyngeal arches in *prdm16* mutants relative to wildtype controls at 24 hpf (Supplemental Fig. 5A-5D), further implicating *prdm16* in pharyngeal arch cell differentiation toward craniofacial derivatives.

**Figure 2.**
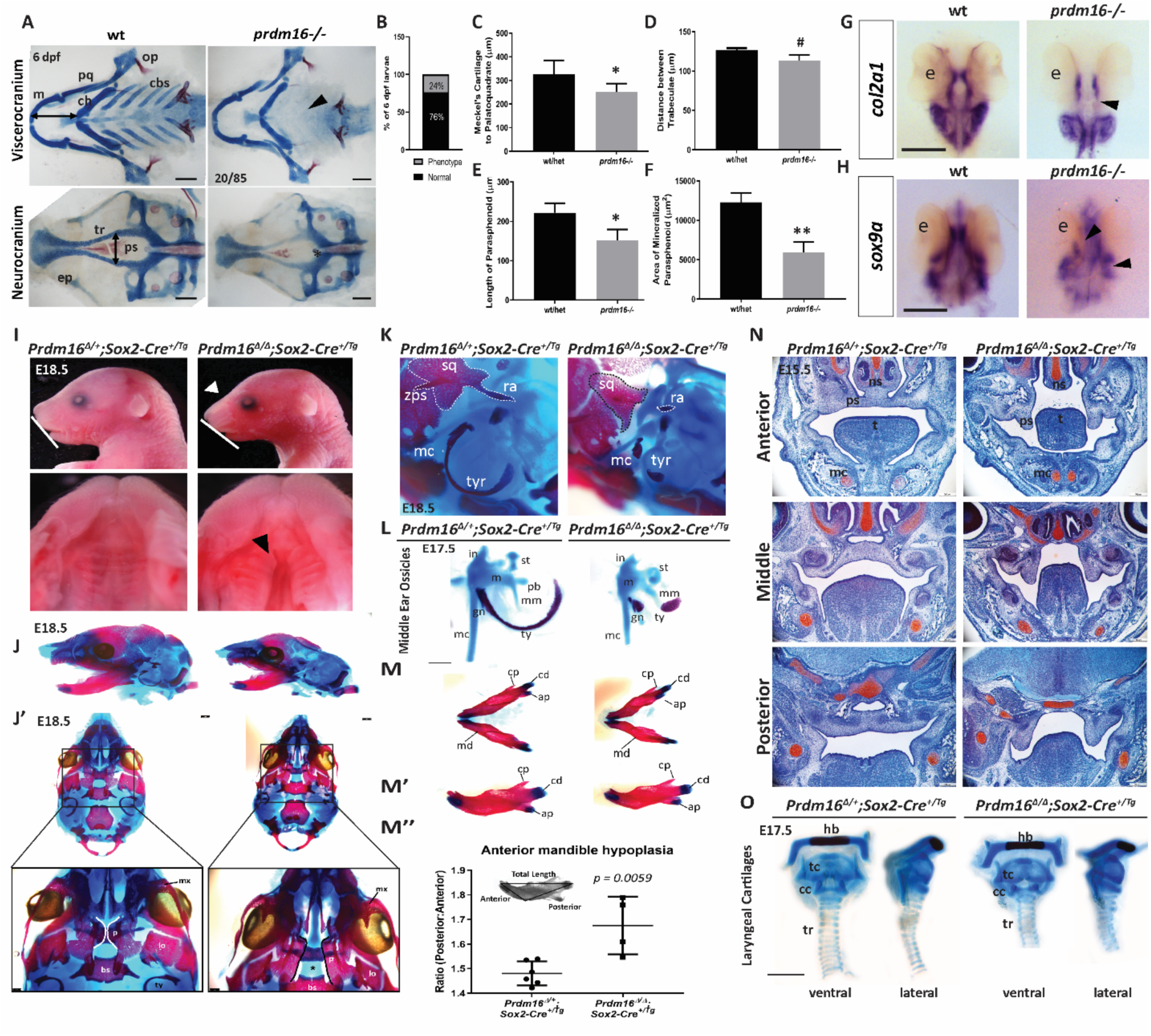
Loss of PRDM16 causes craniofacial defects in zebrafish and mice. (A-E) Wildtype and *prdm16-/-* zebrafish embryos were collected at 6 dpf and stained for Alcian blue and Alizarin red. (A) Images of dissected viscerocranium and neurocranium of wildtype (wt) or *prdm16-/-* embryos. *Prdm16-/-* embryos and their cartilage phenotypes were present in Mendalian ratios (B). (C-F) Quantification of cartilage elements, Meckel’s cartilage to palatoquadrate (C), distance between trabeculae (D), length of parasphenoid tissue (E), and area of mineralized parasphenoid (F). ceratobranchial arches (cbs), ceratohyal (ch), Meckel’s cartilage (m), palatoquadrate (pq), parasphenoid (ps), trabeculae (tr), (n = 6 for each group measured). Scale bar, 100 µm. (G-H) Ventral views of *in situ hybridization* for *col2a1* (G) and *sox9a* (H) in *prdm16* mutants or wildtype controls at 48 hpf. e, eye. Black arrow heads denote areas of reduced expression of these markers in the developing cartilaginous structures. Scale bar, 250 µm. (I-O) *Prdm16^fl/fl^* females were bred for timed matings to *Prdm16^Δ/+^;Sox2^+/Tg^* males and embryos were collected at the indicated timepoints. (I) Lateral images of the head show snout defects (white arrowhead) and hypoplasia of the mandible (white line) in *Prdm16* mutant animals at E18.5 (top). Dissected palates reveal clefting in a subset of *Prdm16* mutants (black arrowheads). (J-M) Alcian blue and Alizarin red stained whole mounts of control and mutants at E18.5. (J) Lateral views of the cranial skeleton. (J’) Ventral views of the cranial skeleton with the mandible removed with higher magnification images of the palantine bone shown below (black star indicates clefting in mutant). Scale bars, 1 mm. (K) High magnification lateral views of the middle ear and squamosal bone structures. (L) Middle ear ossicles were dissected and removed from control or *Prdm16* mutant mice to reveal severe hypoplasia of the malleus, incus and tympanic ring in mutant animals. Scale bar, 0.5 mm. (M-M’’) Mandibles were dissected and removed from control or mutant animals. (M’) Lateral views of the right half of the mandible. (M’’) Quantification of anterior mandible hypoplasia by measuring the ratio between the anterior and posterior portions of the mandible (P:A). (N) Safranin O and Fast Green stained coronal sections through the anterior-posterior axis of the developing palate at E15.5 in control or mutant mice. Scale bars, 250 μm. (O) Dissected and removed laryngeal cartilages from control and *Prdm16* mutant embryos at E17.5 Abbreviations: angular process (ap), basisphenoid (bs), condylar process (cd), coronoid process (cp), cricoid cartilage (cc), gonial (gn), hyoid bone (hb), incus (i), lamina obturans (lo), malleus (m), mandible (md), manubrium of the malleus (mm), maxilla (mx), Meckel’s cartilage (mc), nasal septum (ns), palantine (p), palatal shelf (ps), processus brevis (pb), retroarticular process of the squamosal bone (ra), stapes (st), squamosal bone (sq), thyroid cartilage (tc), tongue (t), tracheal rings (tr), tympanic ring (tyr), zygomatic process of the squamosal bone (zps). * p ≤ 0.05, ** p ≤ 0.005, # p = 0.189, Student’s *t* test.

To assess *PRDM16*’s role in mammalian development, we conditionally ablated *Prdm16* in the murine epiblast with the *Sox2-Cre* driver. Unlike *Prdm3/Evi1*, loss of *Prdm16* in the embryo proper does not cause mid-gestation lethality. Instead, homozygous *Prdm16^Δ/Δ^;Sox2-Cre^+/Tg^*mutants are present at Mendelian ratios with a variety craniofacial defects (Fig. 2I-2O). We did not thoroughly assess the viability of mutant animals past E18.5 but predict, based on previous studies, homozygous *Prdm16^Δ/Δ^;Sox2-Cre^+/Tg^* mutants would not survive postnatally [35]. Gross anatomical observation of animals at E18.5 showed *Prdm16* homozygous mutants have snout extension defects and anterior mandibular hypoplasia (Fig. 2I). Alcian blue and Alizarin Red stained skeletal preparations confirmed the observed mandibular hypoplasia in *Prdm16* mutants (Fig. 2J-J’, 2M-2M”). The anterior portion of the mandible is significantly reduced in mutants, as quantified by measuring the ratio of the anterior length of the mandible relative to the posterior length (Fig. 2M’-2M”).

Previous work has shown ENU-derived *Prdm16* mutant mice develop a secondary cleft palate [35]. Dissected palates revealed a subset of *Prdm16^Δ/Δ^;Sox2-Cre^+/Tg^* mutant mice develop a complete secondary cleft palate (66%, n = 4 out of 6 mutants) (Fig. 2I, 2J’). Skeletal preparations of E18.5 *Prdm16* mutants showed their palatal shelves do not completely meet at the midline and in the most severe cases, there was a complete failure of the palatal shelves to elevate and fuse at the midline (Fig. 2J’). Altered palate development was further examined histologically with coronal sections through the anterior to posterior axis of *Prdm16^Δ/+^;Sox2-Cre^+/Tg^*control and *Prdm16^Δ/Δ^;Sox2-Cre^+/Tg^* mutant mice (Fig. 2N). Safranin O staining of sections showed that at E15.5, *Prdm16^Δ/+^;Sox2-Cre^+/Tg^*control mice have palatal shelves that elevated and began fusing along the anterior to posterior axis (Fig 2N). A cohort of homozygous *Prdm16-Sox2-Cre* mutant animals displayed palatal shelves that had not elevated (50%, n = 4), which likely corresponds to the subset of *Prdm16^Δ/Δ^;Sox2-Cre^+/Tg^*animals that exhibit a complete secondary cleft palate at later stages of development (Fig. 2I, 2J’, 2N).

Notably, all *Prdm16^Δ/Δ^;Sox2-Cre^+/Tg^*embryos develop middle ear defects with severe hypoplasia of the tympanic rings, abnormal formation of the gonium and hypoplasia of the incus and mallus (100%, n = 7) (Fig. 2K-2L). Skeletal preparations also showed that while the squamosal bone formed, it was smaller and misshapen compared to control mice (Fig. 2K). In addition, the zygomatic process of the squamosal bone that extends anteriorly out toward the jugal bone and maxilla was completely missing, as was the retroarticular (retrotympanic) process that extends posteriorly toward the middle ear structures (Fig. 2J’, 2K). Phenotypes in these structures, which are derived from the dorsal mesencephalic neural crest of the first pharyngeal arch [63, 64], exhibited 100% penetrance (Fig. 2J). Hypoplasia of the tympanic rings and middle ear structures in *Prdm16^Δ/Δ^-Sox2-Cre^+/Tg^* mutant mice parallel the hypoplasia of the Meckel’s cartilage and palatoquadrate tissues observed in *prdm16-/-* zebrafish, suggesting PRDM16 function in first and second arch derivatives is conserved across vertebrate species.

In zebrafish, loss of *prdm16* also causes hypoplasia and loss of posterior arch derived structures in the craniofacial skeleton (Fig 2A). To see if PRDM16 has a conserved role in posterior arch derived structures across vertebrates, we assessed posterior pharyngeal arch derived structures in *Prdm16* mutant mice. The laryngeal cartilages, including the thyroid cartilage, cricoid cartilages and tracheal rings are derived from pharyngeal arches 3 and 4. Laryngeal cartilages were dissected from Alcian blue and Alizarin red stained skeletal preparations of heterozygote *Prdm16^Δ/+^;Sox2-Cre^+/Tg^* and *Prdm16^Δ/Δ^;Sox2-Cre^+/Tg^* homozygous animals at E18.5 (Fig. 2O). Interestingly, loss of *Prdm16* only caused subtle differences in the formation of the laryngeal cartilages, mostly an overall reduction in size and moderate compaction of these cartilaginous tissues such as the cricoid cartilage and tracheal rings (Fig. 2O). Together, these results suggest that while PRDM16 function is important for the development of the posterior cartilages in zebrafish, loss of PRDM16 only causes subtle changes to the equivalent posterior arch derivatives in mice.

### *prdm3* and *prdm16*, along with *prdm1a* compensate for each other in zebrafish

Vertebrate genomes contain 17 different PRDM family members that generally share structural similarity with a combination of conserved N-terminal PR domain, as well as a varying number of zinc fingers. The zebrafish genome appears to only have duplicated *prdm8* (*prdm8a*, *prdm8b*) and *prdm1* (*prdm1a*, *prdm1b*, *prdm1c*), all other zebrafish *prdm* genes are not duplicated. PRDM3 and PRDM16 are very closely related, sharing 63% nucleotide and 56% amino acid identity, as well as similar domain structures [65]. In addition, zebrafish morphants for *prdm3* and *prdm16* show they share redundant roles in zebrafish craniofacial development and they genetically interact with another member of the PRDM family, *prdm1a* to facilitate craniofacial skeletal formation [36]. Finally, work from our lab and others in zebrafish and mice shows PRDM1 (Blimp1) also is essential for craniofacial development, as *prdm1a* is required for the formation of posterior craniofacial structures and cranial neural crest-derived glands [51, 66, 67].

As such, we asked whether the modest craniofacial phenotypes observed in *prdm3* and *prdm16* single zebrafish mutants was the result of functional redundancy and thus genetic compensation between *prdm3*, *prdm16* and *prdm1a*. To test for genetic compensation, the mRNA expression of each of these *prdms* (*prdm1a*, *prdm3* and *prdm16*) was assessed by qPCR on RNA extracted from whole heads isolated from *prdm3-/-* and *prdm16-/-* mutants at 48 hpf (Fig. 3A-3C). In support of genetic compensation, *prdm16* expression was elevated in *prdm3-/-* mutants (Fig. 3B). Interestingly, however, there was a reduction in *prdm3* expression in *prdm16-/-* mutants (Fig. 3A). This could explain the slight differences in the phenotypes between *prdm3-/-* and *prdm16-/-*, with *prdm16-/-* embryos exhibiting moderately more severe posterior ceratobranchial cartilage phenotypes compared to *prdm3* mutants (Fig. 2A). On the other hand, *prdm1a* mRNA was increased in both *prdm3-/-* and *prdm16-/-* embryos (Fig. 3C). This was further confirmed by *in situ hybridization* for *prdm1a* performed on 24 hpf *prdm3* and *prdm16* mutant embryos (Fig. 3D). In these embryos, *prdm1a* expression was not only increased, but the domains of expression were expanded (Fig. 3D).

**Figure 3.**
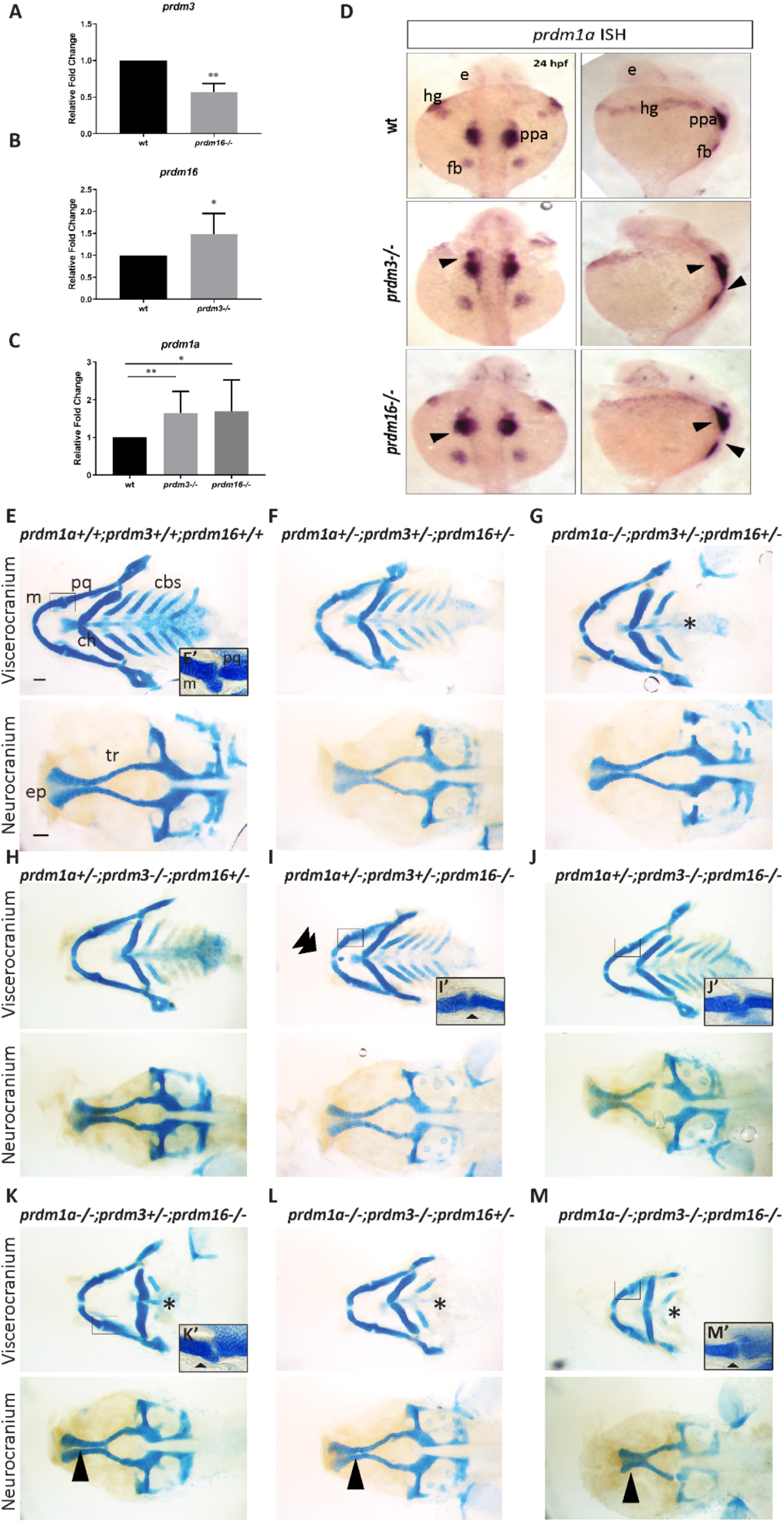
*prdm1a, prdm3*, and *prdm16* genetically compensate for each other in zebrafish craniofacial development. (A-C) RNA was isolated from the heads of *prdm3-/-*, *prdm16-/-* or wildtype embryos at 48 hpf. RT-qPCR was performed for *prdm3* (A), *prdm16* (B) or *prdm1a* (C). 5-10 genotyped heads were pooled for RNA. (n = 3 replicate experiments). (D) Dorsal and lateral views of *in situ* hybridization for *prdm1a* in *prdm3* and *prdm16* mutants at 24 hpf. (E-M) *prdm1a+/-;prdm3+/-;prdm16+/-* triple heterozygous fish were generated and intercrossed. Embryos were collected and stained at 4 dpf with Alcian blue. Shown are dissected and mounted viscerocranium and neurocranium tissues from different allelic combinatorial mutants indicated. Selected inserts (E’, I’-K’, M’) show high magnification images of the jaw joints. Double arrowhead indicates ectopic cartilage, black star indicates loss of posterior ceratobranchial arches, single short black arrowhead indicates abnormal jaw joint, single long black arrowhead indicates clefting in the ethmoid plate. Abbreviations: ceratobranchial arches (cbs), ceratohyal (ch), ethmoid plate (ep), eye (e), fin bud (fb), hatching gland (hg), Meckel’s cartilage (m), palatoquadrate (pq), posterior pharyngeal arches (ppa), trabeculae (tr). Scale bar, 100 µm. * p ≤0.05, ** p ≤ 0.005, Student’s *t* test.

To further understand the genetic compensation between *prdm1a*, *prdm3*, and *prdm16*, we generated *prdm3+/-;prdm16+/-* double heterozygotes and bred these into the established *prdm1a+/-* mutant line, *narrowminded* (*nrd*) [68] to generate *prdm1a+/-;prdm3+/-;prdm16+/-* triple heterozygotes. These animals survived to adulthood and were fertile so they were used for intercross breedings. Only one *prdm1a-/-;prdm3-/-;prdm16-/-* triple mutant (out of 5 intercross breedings) survived to 4 dpf, the earliest stage at which craniofacial cartilage development was assessed (Fig. 3M). In support of our hypothesis of genetic compensation by *prdm1*, Alcian blue staining of 4 dpf compound mutant larvae revealed more severe craniofacial defects (Fig. 3E-3M). Dissected skeletal preparations showed complete loss of the posterior ceratobranchial cartilage elements in all compound mutants harboring homozygosity for *prdm1a*, a phenotype consistent with *prdm1a* single mutants (Fig. 3G, 3K, 3L, 3M). *prdm1a-/-;prdm3-/-;prdm16+/-* and *prdm1a-/-;prdm3-/-;prdm16-/-* showed increased severity of the hypoplasia of the anterior viscerocranium cartilages, relative to single mutants alone, including the Meckel’s cartilage and palatoquadrate (Fig. 3L-3M). Additionally, some combinatorial mutants, particularly those that were homozygous for *prdm16,* had either loss or severe hypoplasia of articular process of the Meckel’s cartilage or this process was fused to the palatoquadrate leading to an abnormal jaw joint (Fig. 3I’, 3J’, 3K’, 3M’). This phenotype correlates to the middle ear defects observed in *Prdm16^Δ/Δ^;Sox2-Cre^+/Tg^*mutant mice, as the middle ear of tetrapods is evolutionarily derived from the jaw joint of non-mammalian vertebrates such as the zebrafish [69–77]. Notably, near or complete homozygosity for all three alleles led to clefting in the ethmoid plate (Fig. 3K-3M), also mirroring the cleft palate phenotype we observed in *Prdm16^Δ/Δ^-Sox2-Cre^+/Tg^*mutant mice. Together, these results show *prdm3* and *prdm16* likely act redundantly during craniofacial development. These results further suggest *prdm3* and *prdm16* genetically interact with *prdm1a* to drive proper formation of the craniofacial skeleton. Loss of either *prdm3* or *prdm16* causes only minor craniofacial defects due to compensation between each other and with *prdm1a*. Lastly, these results demonstrate an evolutionarily conserved functional role for Prdm16 across mammalian and non-mammalian vertebrates in the formation and maintenance of the mammalian middle ear and the non-mammalian jaw joint.

### Loss of *prdm3* and *prdm16* in zebrafish and mice decreases trimethylation of histone targets

To begin to disentangle the mechanism by which PRDMs function, we assessed methylation of the chromatin marks that have been identified as the primary substrates for PRDM3 and PRDM16. PRDM3 and PRDM16 act to methylate H3K9 and repress gene expression [21] and in more recent studies have also been shown to methylate H3K4 to activate gene expression in certain cellular contexts [22, 78, 79]. As expected, methylation of the target residue, H3K9 was significantly reduced by 31% in *prdm3-/-* and 58% in *prdm16-/-* whole zebrafish embryo lysates at 48 hours post fertilization (hpf) (Fig. 4A, 4B). Similarly, methylation of the activating mark, histone 3 lysine 4 (H3K4) was reduced by 40% in *prdm3* mutant embryos and 50% in *prdm16* embryos (Fig. 4C, 4D). We next explored the methylation of these two histone marks in the palatal shelves of *Prdm16^Δ/Δ^;Sox2-Cre^+/Tg^*mouse embryos, whereas lethality of *Prdm3^Δ/Δ^;Sox2-Cre^+/Tg^*mice did not allow this assessment. In heterozygote *Prdm16^Δ/Δ^;Sox2-Cre^+/Tg^* control mice, immunofluorescence staining on coronal sections through the palate showed H3K9me3 was abundant in the palatal mesenchyme cells at E15.5, the stage at which the palatal shelves have elevated and fused (Fig. 4E, 4G). Methylation of H3K9 was significantly reduced in the palatal shelves of *Prdm16^Δ/Δ^;Sox2-Cre^+/Tg^*mutant mice that had failed to elevate (Fig. 4E, 4G). This drastic difference in H3K9me3 was quantified by measuring total cell fluorescence in different fields of view and corrected for area and background fluorescence (CTCF) (Fig. 4E). Conversely, we did not see a change in the activating mark, H3K4me3 between homozygous mutants and control animals at this stage (Fig. 4H, 4F). The difference between complete loss of H3K9me3 and no change in H3K4me3 in the context of mammalian craniofacial and palate development, suggests PRDM16 acts more in favor of gene repression rather than as an activator of gene expression. The comparable decreases in H3K9me3 across both zebrafish and *Prdm16^Δ/Δ^-Sox2-Cre^+/Tg^*mutant mice suggest Prdm3 and Prdm16 are regulating craniofacial development through broad epigenetic mechanisms associated with gene repression of different developmental programs. The difference in H3K4me3 between zebrafish and mice indicates the potential for divergent mechanisms of craniofacial development across these vertebrates. Specifically, these results suggest Prdm3 and Prdm16 may act not only as repressors but as activators during zebrafish craniofacial development. This dual role of gene activation and repression through modification of these chromatin marks may not be conserved across mammalian craniofacial development, at least at the level of H3K4 methylation. Taken together, PRDM3 and PRDM16 control the chromatin landscape to mediate proper repression and/or activation of specific developmental gene regulatory networks during craniofacial development in mammalian and non-mammalian vertebrates.

**Figure 4.**
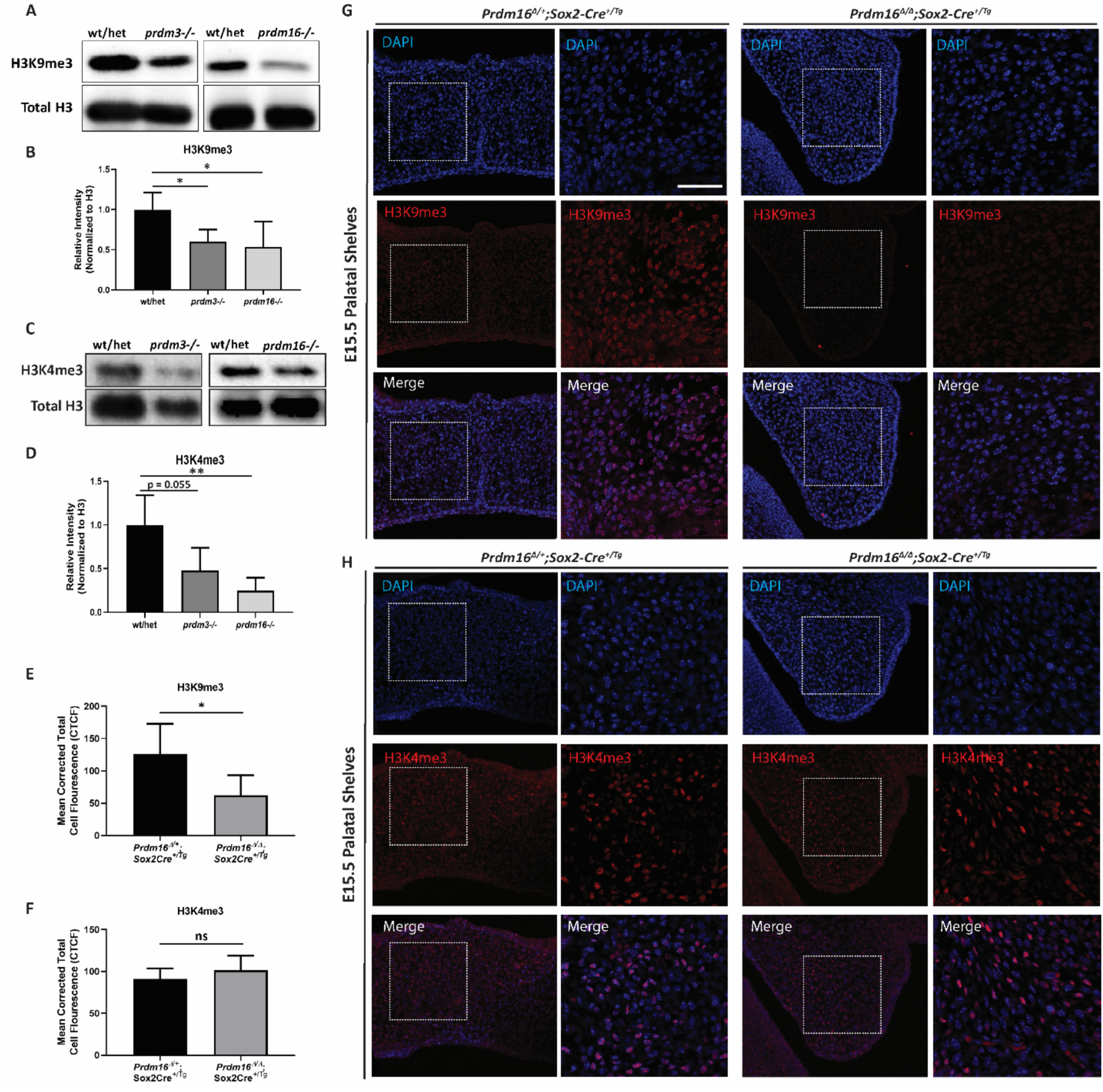
Loss of *prdm3* and *prdm16* decreases trimethylation of histone H3 substrates. (A-D) Wildtype, *prdm3-/-* and *prdm16-/-* single zebrafish mutants were pooled at 48 hpf and harvested for whole embryo protein lysates. (A) Proteins were resolved by SDS-PAGE and western blotting for H3K9me3 (A), H3K4me3 (C) and total H3 was performed. (B, D) Quantification of band intensity of the western blots in (A and C) for H3K9me3 (B) or H3K4me3 (D) (n = 3 different experiments). Relative intensity was normalized to total H3 for each sample for all replicate experiments. All mutant samples were then normalized to the average of normalized intensity from the controls and averaged. (E-H) Immunofluorescence for H3K9me3 (G) or H3K4me3 (H) was performed on coronal sections of the palatal shelves in E15.5 control or *Prdm16^Δ/Δ^;Sox2-Cre^+/Tg^* mice. Shown are lower magnification (20x objective) and higher magnification (63x objective) of the indicated areas. (E-F) Quantification of the mean corrected total cell fluorescence (CTCF) from 5 different fields of view averaged across n = 2-3 embryos per group. Scale bar 50 µm. * p ≤ 0.05, ** p ≤0.005, ns, not significant, Student’s *t* test.

## Discussion

This study elucidates the essential functions of PRDM3 and PRDM16 in proper craniofacial skeletal development in zebrafish and mice. We show that loss of these two Prdm genes in zebrafish causes anterior and posterior cartilage defects in the viscerocranium and neurocranium in the developing craniofacial skeleton. In addition, *prdm3* and *prdm16* together with *prdm1a* can compensate for each other and that combinatorial loss of all three of these alleles causes drastically more severe craniofacial phenotypes. Consistent with previous studies, we demonstrate that while genetically ablating *Prdm3/Evi1* in the murine epiblast causes mid-gestation lethality, loss of *Prdm16* in the early embryo is not lethal and instead causes anterior and posterior cartilage and bone defects. Finally, we describe both conserved and divergent functionality for Prdm3 and Prdm16 across mice and zebrafish through modulation of histone marks and dynamically controlling the chromatin landscape to activate or repress gene regulatory networks required for craniofacial development.

Of the 17 PRDM histone methyltransferase family members, PRDM3 and PRDM16 are the most similar structurally at the amino acid and protein structure level, which strongly suggests functional redundancy between the two. In our zebrafish mutants, we observed compensation between not only *prdm16* in *prdm3* mutants, but *prdm1a* in both *prdm3* and *prdm16* single mutants emphasizing the shared functional roles of these factors. Furthermore, the most severe craniofacial phenotypes only develop upon combinatorial loss of all three alleles. These findings implicate functional redundancy across these PRDMs in the context of craniofacial development. While *prdm3*, *prdm16* and *prdm1a* have very similar expression patterns during early zebrafish development, with overlapping expression in the pharyngeal arches and hindbrain, they do have differences between their developmental expression patterns. For example, only *prdm16* is strongly expressed in the olfactory placodes while only *prdm3* expression is detected in the ventral diencephalon and the tegmentum [36]. Variations in expression patterns suggest that while these Prdms can compensate for each other, they may also have different functions depending on cellular context and developmental time. It is likely that loss of any of the three PRDM alleles initiates compensatory mechanisms from the other paralogs, allowing for retention of PRDM functionality. It is also possible that within the mutant background, the normal repressive function may instead lead to activation with results in misregulation of gene expression. Variations in functional redundancy between *prdm3* and *prdm16* is also emphasized with the differences in severity between single and compound mutants. Loss of *prdm16* alone appears to be more drastic than *prdm3* alone. Additionally, *prdm3* does not compensate in *prdm16* mutants to the extent that *prdm16* compensates in *prdm3* mutants suggesting *prdm16* maybe more important than *prdm3* within this cellular context of development. Future experiments utilizing non-biased whole transcriptomics are required to better distinguish the similarities and differences in functionality between Prdm3 and Prdm16 during craniofacial development.

While Prdm3 and Prdm16 are among the few PRDMs that exhibit intrinsic methyltransferase activity, they also form complexes with other factors including transcription factors as well as histone modifying enzymes to control gene expression. Prdm1, on the otherhand, has been shown to regulate gene expression *in vivo* through interaction with protein complexes. Interestingly, while there are a couple proteins Prdm3 and Prdm16 are known to both bind (e.g. CtBP, Smad3) [29, 39, 41, 80], overall, there is little overlap between the currently known binding partners across all three Prdms. The context of the protein complex(es) most likely determines whether the Prdm proteins establish a protein-protein or DNA-protein interaction to facilitate gene activation or repression. The diversity between binding partners could also explain some variation in phenotypic severity across compound mutants as well as further define differences in functionality across these highly similar Prdm family members. In addition to utilizing whole trancriptomics to better understand the differences and similarities between Prdm3- and Prdm16-dependent trancripional changes, the identification of the protein binding partners that drive activation and repression during craniofacial development will establish the molecular mechanism of compensation and specificity of Prdm3 and Prdm16.

We describe potential conserved roles of PRDM3 and PRDM16 from zebrafish to mice, mostly through the similarities observed in the craniofacial skeletons with loss of function of these Prdms in both species. While the mid-gestation lethality due to genetically ablating *Prdm3/Evi1* in the early developing embryo hinders understanding the function of *Prdm3* in craniofacial development, our early cartilage whole mounts suggest that loss of *Prdm3* leads to hypoplasia of the Meckel’s cartilage. Along with the phenotypes observed in *Prdm16^Δ/Δ^-Sox2-Cre^+/Tg^* mutant mice (i.e. anterior mandibular hypoplasia, secondary cleft palate, tympanic ring/middle ear hypoplasia) and those observed in *prdm3* and *prdm16* single and combinatorial mutant zebrafish, structures derived from the first and second pharyngeal arches are the most sensitive to Prdm loss in both vertebrates. Furthermore, loss of *Prdm16* in mouse resulted in severe hypoplasia of the middle and outer ear structures (malleus, incus, tympanic rings). The mammalian inner ear is evolved from the non-mammalian vertebrate jaw joint, where the mammalian malleus and incus correspond to the articular process of the proximal end of the Meckel’s cartilage and the palatoquadrate, respectively [69–77]. In compound *prdm1a;prdm3;prdm16* zebrafish mutants, the articular process of the Meckel’s cartilage is either lost, hypoplastic, or partially fused to the palatoquadrate. Interestingly, one of the more striking phenotypes in the zebrafish mutants (compounds with homozygosity for *prdm1a*, but also *prdm16* single mutants) is the loss of the posterior ceratobranchial arches. We did not observe major changes in the posterior derived laryngeal cartilages in *Prdm16^Δ/Δ^-Sox2-Cre^+/Tg^*mutant mice, suggesting divergent roles for PRDM16 in the posterior arches. Nevertheless, the similarities between the non-mammalian jaw and the mammalian middle ear implicate a highly conserved role for these Prdms, and in particular, Prdm16, in the establishment and maintenance of these particular craniofacial structures across vertebrates. While mechanisms that allow for these Prdms to mediate the proper development and function of these craniofacial structures remains largely unknown, our data suggest that this function maybe evolutionarily conserved in some craniofacial tissues. More experiments are required to establish a cell autonomous role for these Prdms specifically in the neural crest and the formation of the craniofacial skeleton. In addition, we observe functional redundancy between Prdm3 and Prdm16 in our zebrafish mutants. It is possible functional redundancy for *Prdm1*, *Prdm3*, *Prdm16* exists in mammals. While assessing functional redundancy is complicated by the mid-gestation lethality associated with other roles of *Prdm3/Evi1* in vasculogenesis, future experiments will involve evaluation of this redundancy specifically in the neural crest lineage.

Finally, we have established a necessary and conserved role for Prdm1a, Prdm3, and Prdm16 in the development of the craniofacial skeleton in zebrafish and mice by modulating chromatin state. As such, the phenotypes in our animal models could be indirect secondary effects associated with broadly altering the chromatin landscape. We observed changes not only associated with the canonical repressive histone mark (H3K9me3) typically mediated by these PRDMs, but we also showed alterations in activating histone marks (H3K4me3) [21, 22]. These findings support a mechanism by which these factors act dynamically during development to not only repress certain gene regulatory networks but activate other networks depending on the time and cellular context. Recent studies have implicated Prdm16 in activation of gene expression by associating with H3K27ac, and directly binding active enhancers in the context of cortical neuron development [23]. Future experiments are needed to identify the direct targets of these broad methyltransferases and assess global changes in the chromatin landscape with loss of these factors in order to establish specific regulatory roles (both active and repressive) for these Prdms spatially and temporally in the context of neural crest development and the formation of the craniofacial skeleton.

## Conclusions

In summary, we have identified Prdm3 and Prdm16, with Prdm1 as crucial mediators of neural crest differentiation toward craniofacial skeleton elements. We show that loss of these factors does not affect neural crest migration, but rather impedes proper differentiation and subsequent formation of the cartilaginous and bone structures required for craniofacial development. We also show that these methyltransferases share functional redundancy in zebrafish and importantly that they share evolutionarily conserved functions across mammalian and non-mammalian vertebrates. Understanding the specific functional roles of these dynamic factors during development is crucial towards identifying how alterations to these regulatory networks drive congenital birth defects including mandibular hypoplasia, cleft palate and abnormal middle ear development.

## Acknowledgments

We thank our collaborators Dr. Lee Niswander and Sofia Pezoa for our ongoing collaboration, continuous feedback on this project and for insightful comments on the manuscript; Dr. Mineo Kurokawa for the Prdm3^fl/fl^ mouse line and Jason Willams, Kristi LaMonica and Sofia Pezoa for help maintaining the mouse lines; and David Clouther for mouse craniofacial antomy consultation. We thank the Denver zebrafish community and the zebrafish facility staff for excellent fish care.

## Funding

This work is supported by NIH/NIDCR NRSA 1 F32 DE029099-01 postdoctoral fellowship to L.C.S., ACS postdoctoral fellowship to R.S., NIH/NIDCR R01DE024034 to K.B.A. and NIH NINDS P30NS048154 to the UC Denver zebrafish core facility.

---

**Supplemental Figure 1.**
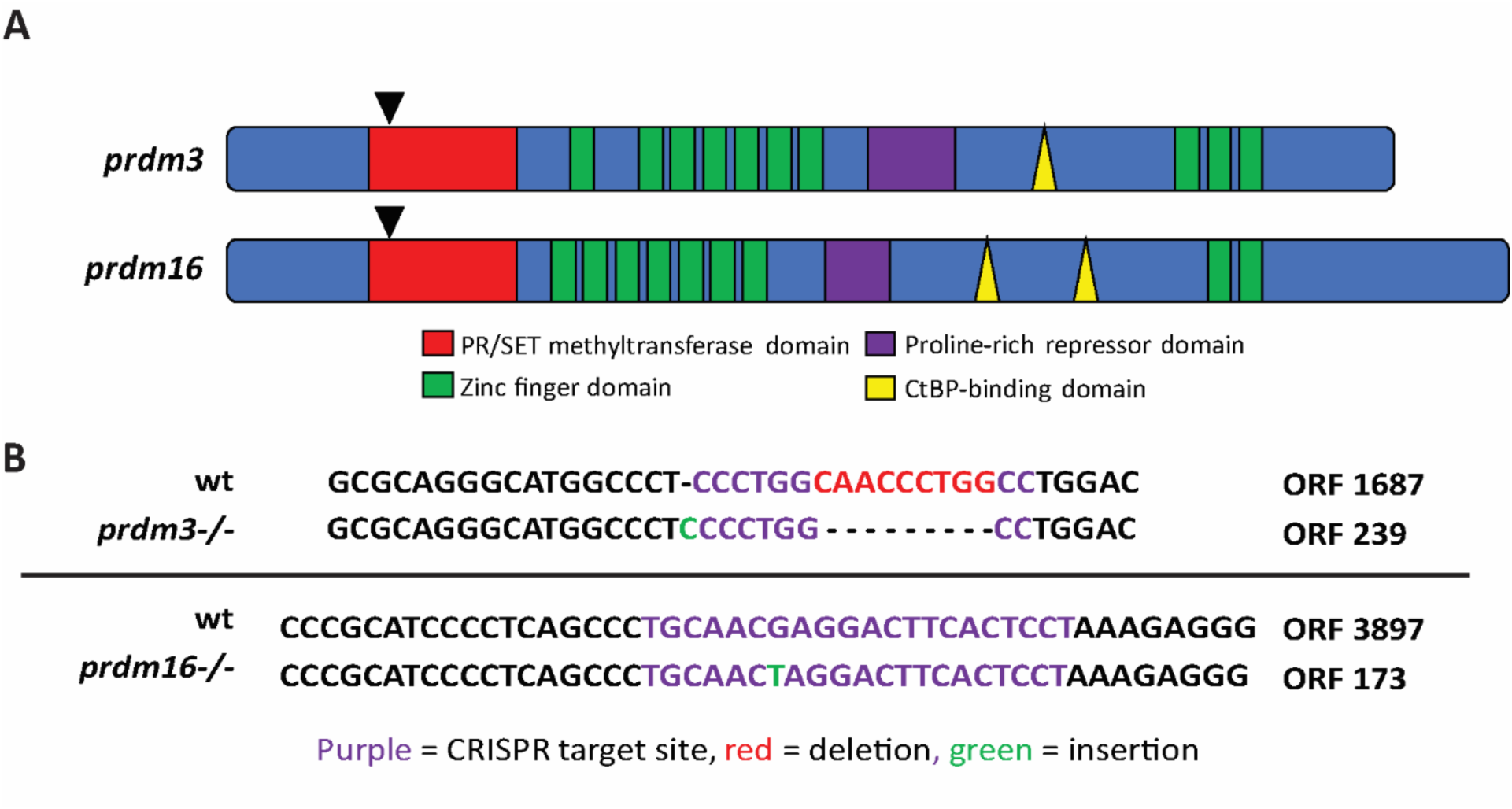
CRISPR/Cas9 mediated genome editing at the prdm3^CO1005^ and prdm16^CO1006^ loci creates frameshift mutations resulting in truncated proteins. (A) Schematic of the structures of *prdm3* and *prdm16,* highlighting the specific domains, including the PR/SET methyltransferase domain (red), zinc finger domains (green). Black arrowheads indicate the location of CRISPR/Cas9 mediated mutation for each gene. (B) CRISPR/Cas9 directed genome editing created an indel (green) and a 9 base pair deletion (red) in the CRISPR target site (purple) in *prdm3* and an indel (green) in the *prmd16* CRISPR target site, both resulting in frameshifts and truncated proteins.

**Supplemental Figure 2.**
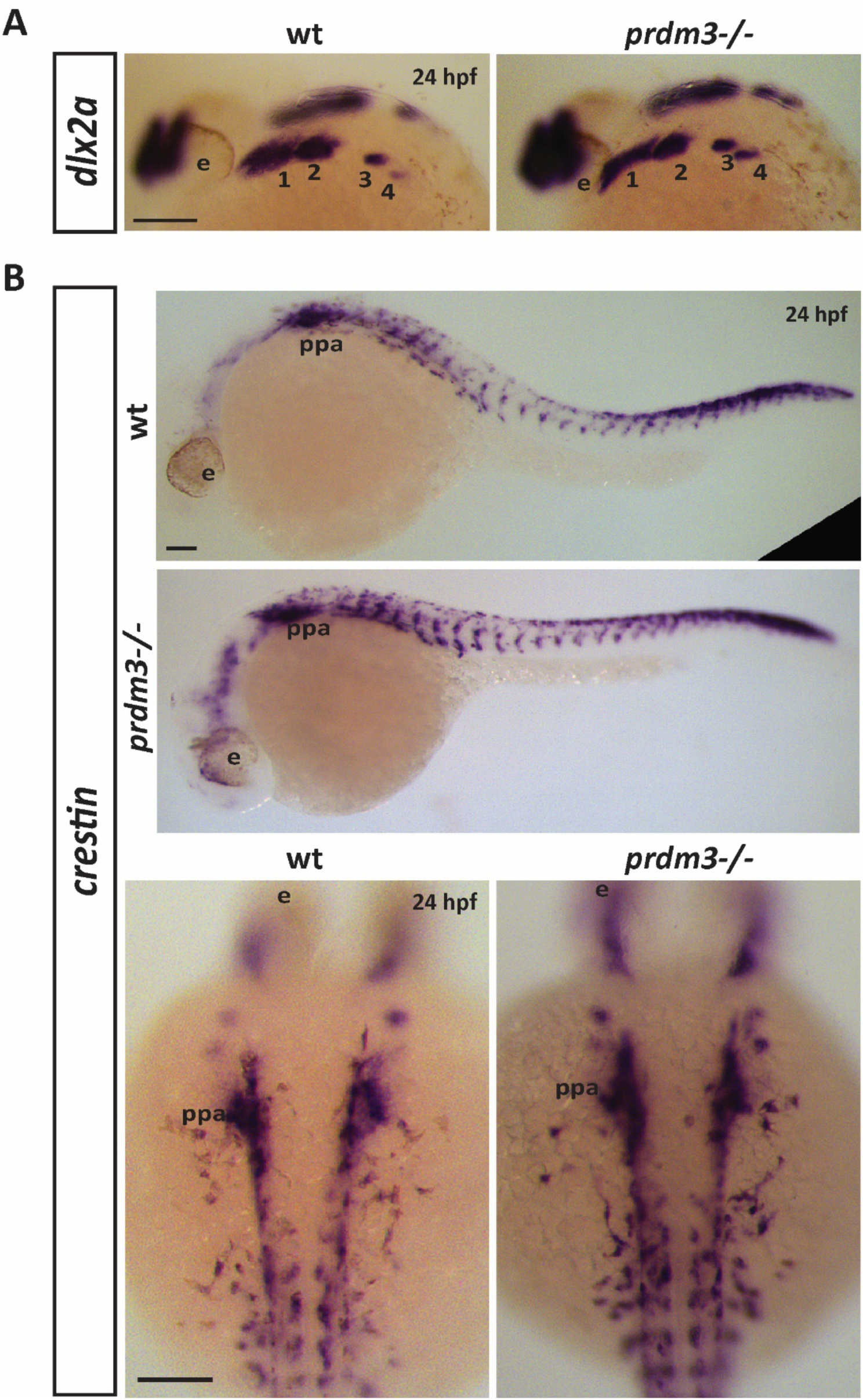
Neural crest markers, *dlx2a* and *crestin*, are unchanged in the pharyngeal arches with loss of *prdm3* in zebrafish. (A-B) Wildtype or *prdm3-/-* mutant embryos were collected and *in situ hybridization* was performed for *dlx2a* (A) or *crestin* (B) at 24 hpf. Shown are lateral views of *dlx2a* expression in the pharyngeal arches (1-4) (A) and lateral views of *crestin* across the whole body to show neural crest migration is unaffected in *prdm3* mutant zebrafish. Higher magnification, dorsal views of *crestin* expression is also shown in (B). Scale bars, 100 µm. Abbreviations: eye (e), 1-4 (pharyngeal arches 1-4), posterior pharyngeal arches (ppa).

**Supplemental Figure 3.**
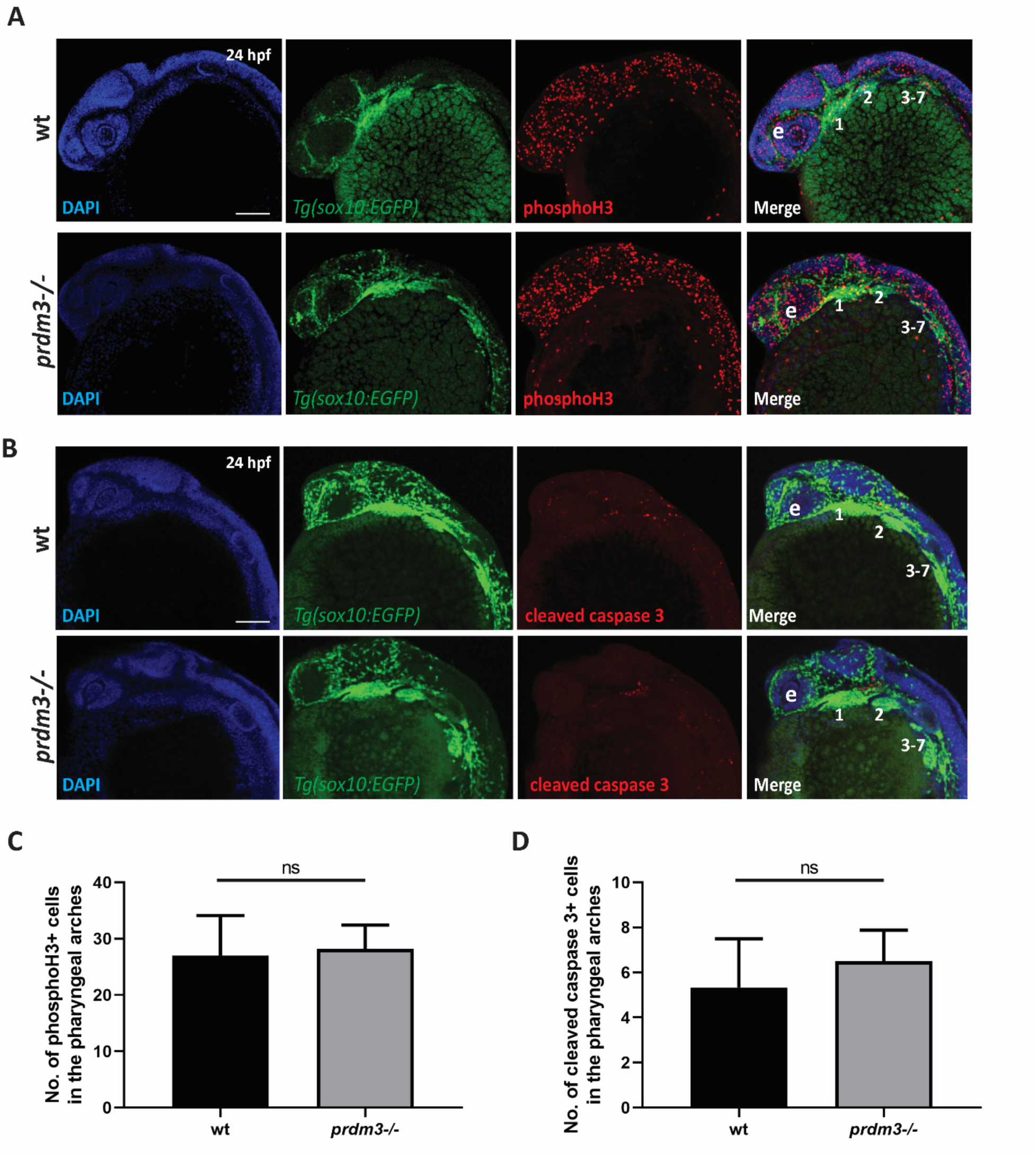
Loss of *prdm3* does not lead to changes in pharyngeal arch cell proliferation or cell death. (A-D) Wildtype or *prdm3-/-* zebrafish embryos crossed into the *Tg(sox10:EGFP)* transgenic line were collected at 24 hpf and whole mount immunofluorescence was performed for phosphorylated histone H3 (A) or cleaved caspase 3 (B) to assess cell proliferation or apoptosis in the pharyngeal arches, respectively. Shown are lateral max projection images of the pharyngeal arches 1-7. (C-D) Quantification of the number of phosphorylated histone H3 positive cells (yellow) (C) or cleaved caspase 3 positive cells (yellow) (D) within the pharyngeal arches (n = 6 embryos for each group). Abbreviations: eye (e), pharyngeal arches (1-7). Scale bar, 100 µm. ns, not significant, Student’s *t* test.

**Supplemental Figure 4.**
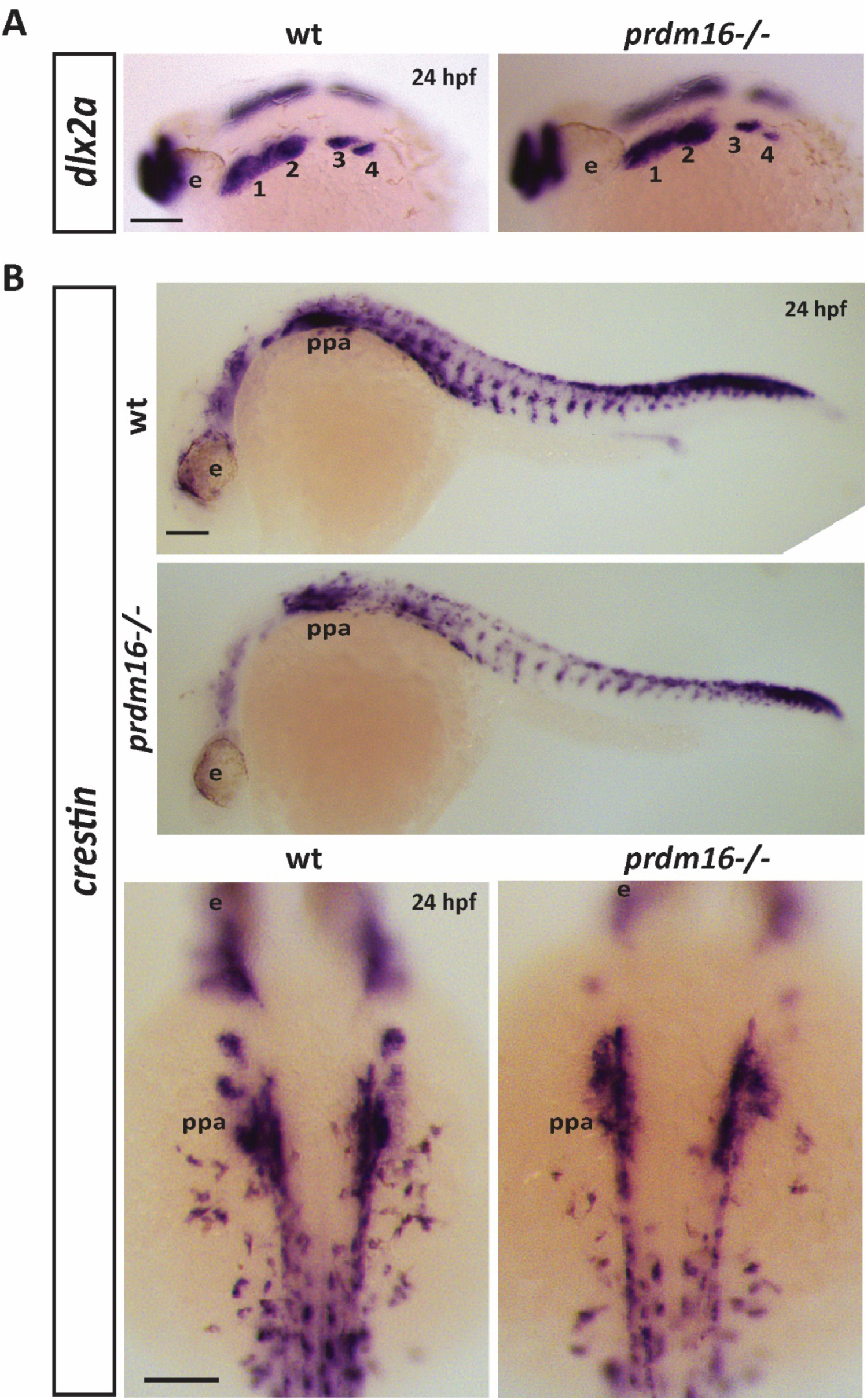
Loss of *prdm16* causes a subtle decrease in expression of neural crest markers *dlx2a* or *crestin* in the pharyngeal arches. (A-B) Wildtype or *prdm16-/-* mutant embryos were collected and *in situ hybridization* for *dlx2a* (A) or *crestin* (B) was performed at 24 hpf. Shown are lateral views of *dlx2a* expression in the pharyngeal arches (1-4) (A) and lateral views of *crestin* across the whole body to show neural crest migration is unchanged in *prdm16* mutant zebrafish. Higher magnification, dorsal views of *crestin* expression is also shown in (B). Scale bars, 100 µm. Abbreviations: eye (e), 1-4 (pharyngeal arches 1-4), posterior pharyngeal arches (ppa).

**Supplemental Figure 5.**
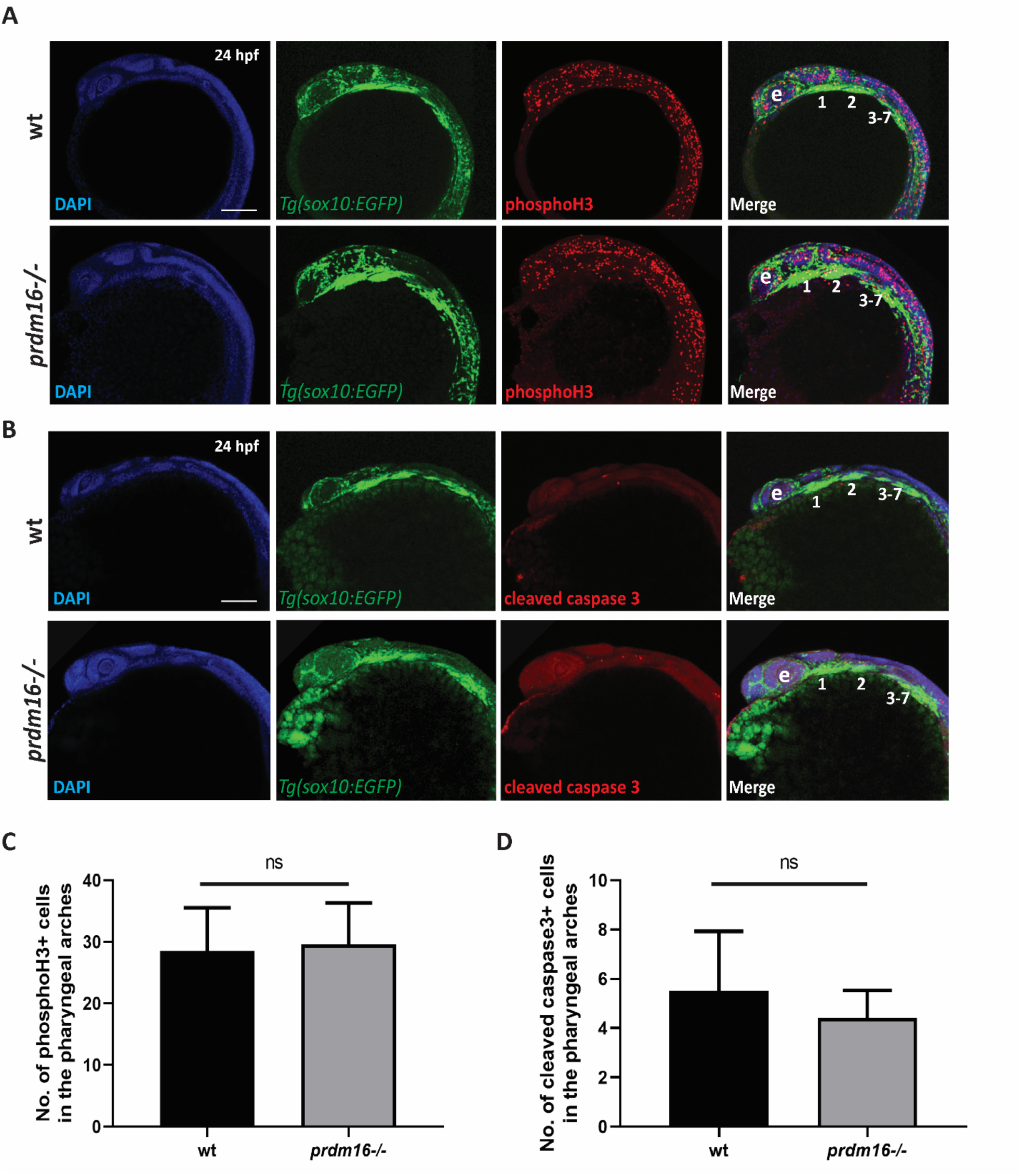
Cell proliferation and cell death are not affected in the pharyngeal arches with loss of *prdm16*. (A-D) Wildtype or *prdm16-/-* zebrafish embryos crossed into the *Tg(sox10:EGFP)* transgenic line were collected at 24 hpf and whole mount immunofluorescence was performed for phosphorylated histone H3 (A) or cleaved caspase 3 (B) to assess cell proliferation or apoptosis in the pharyngeal arches, respectively. Shown are lateral max projection images of the pharyngeal arches 1-7. (C-D) Quantification of the number of phosphorylated histone H3 positive cells (yellow) (C) or cleaved caspase 3 positive cells (yellow) (D) within the pharyngeal arches (n = 6 embryos for each group). Abbreviations: eye (e), pharyngeal arches (1-7). Scale bar 100 µm. ns, not significant, Student’s *t* test.

